# The relationship between spatial configuration and functional connectivity of brain regions

**DOI:** 10.1101/210195

**Authors:** Janine D. Bijsterbosch, Mark W. Woolrich, Matthew F. Glasser, Emma C. Robinson, Christian F. Beckmann, David C. Van Essen, Samuel J. Harrison, Stephen M. Smith

## Abstract

Brain connectivity is often considered in terms of the communication between functionally distinct brain regions. Many studies have investigated the extent to which patterns of coupling strength between multiple neural populations relates to behavior. For example, studies have used "functional connectivity fingerprints" to characterise individuals' brain activity. Here, we investigate the extent to which the exact spatial arrangement of cortical regions interacts with measures of brain connectivity. We find that the shape and exact location of brain regions interact strongly with the modelling of brain connectivity, and present evidence that the spatial arrangement of functional regions is strongly predictive of non-imaging measures of behaviour and lifestyle. We believe that, in many cases, cross-subject variations in the spatial configuration of functional brain regions are being interpreted as changes in functional connectivity. Therefore, a better understanding of these effects is important when interpreting the relationship between functional imaging data and cognitive traits.

## Introduction

The organisation of the human brain into large-scale functional networks has been investigated extensively over the past two decades using resting state functional magnetic resonance imaging (rfMRI). Spontaneous fluctuations in distinct brain regions (as measured with rfMRI) show temporal correlations with each other, revealing complex patterns of functional connectivity (FC) (Biswal, Yetkin, Haughton, & Hyde, 1995; Friston, 1994, 2011). Extensive connectivity between cortical areas and with subcortical brain regions has long been considered a core feature of brain anatomy and function (Crick & Jones, 1993), and dysfunctional coupling is associated with a variety of neurological and psychiatric disorders including schizophrenia, depression, and Alzheimer’s disease (Castellanos, Di Martino, Craddock, Mehta, & Milham, 2013). Given the great potential neuroscientific and clinical value of rfMRI, it is important to determine which aspects of rfMRI data most sensitively and interpretably reflect trait variability across subjects. At a neural level, potential sources of meaningful cross-subject variability include: i) the strength of the functional coupling (i.e., interactions) between two different neural populations (‘*coupling’*), and ii) the spatial configuration and organisation of functional regions (‘*topography’*). In this study, we aim to identify how these key aspect of rfMRI data influence derived measures of functional connectivity and how they relate to interesting trait variability in behaviour and lifestyle across individuals. Our findings reveal variability in the spatial topography of functional regions across subjects, and suggest that this variability is the primary driver of cross-subject trait variability in correlation-based FC measures obtained via group-level rfMRI parcellation approaches. These results have important implications for future rfMRI research, and for the interpretation of FC findings.

A commonly applied approach used to derive FC measures from rfMRI data is to parcellate the brain into a set of functional regions (‘nodes’), and estimate the temporal correlations between pairs of node timeseries (‘edges’) to build a network matrix (Smith, Vidaurre, et al., 2013). This approach has previously been likened to a fingerprint, enabling the unique identification of individuals, and the prediction of behavioural traits such as intelligence (Finn et al., 2015; Passingham, Stephan, & Kötter, 2002). Parcellation methods include the use of anatomical, functional, and multi-modal atlases (Glasser et al., 2016; Tzourio-Mazoyer et al., 2002; Yeo et al., 2011), with functional parcellations often being data driven via techniques such as clustering and independent component analysis (ICA) (Beckmann, DeLuca, Devlin, & Smith, 2005; Craddock, James, Holtzheimer, Hu, & Mayberg, 2012). Data-driven approaches such as ICA have been used to identify consistent large-scale resting state networks (Damoiseaux et al., 2006), and to characterise FC abnormalities in a variety of mental disorders (Littow et al., 2015; Pannekoek et al., 2015). A single parcellation is typically defined at the group level (for any parcellation method), and hence additional steps are required to map a group-level parcellation onto individual subjects’ data in order to obtain subject-specific parcel timeseries and associated connectivity edge estimates. Timeseries derived from hard (binary, non-overlapping) parcellations are often obtained using a simple masking approach (i.e., extracting the averaged BOLD timeseries across all voxels or vertices in a node), whereas ICA parcellations (partially overlapping, soft parcellations that contain continuous weights) are mapped onto single-subject data using dual regression analysis or back projection (Calhoun, Adali, Pearlson, & Pekar, 2001; Filippini et al., 2009). Previous work has shown that, in the presence of spatial variability or inaccurate intersubject alignment, these common methods for mapping group parcellations onto individuals are not able to fully recover accurate subject-specific functional regions, which can severely impact the accuracy of estimated FC edges (Allen, Erhardt, Wei, Eichele, & Calhoun, 2012; Smith et al., 2011).

Recent work has characterised patterns of spatial variability in network topography across subjects (i.e., spatial shape, size and position of functional regions) (Glasser et al., 2016; Gordon, Laumann, Adeyemo, Gilmore, et al., 2016; Gordon, Laumann, Adeyemo, & Petersen, 2015; Laumann et al., 2015; Swaroop Guntupalli & Haxby, 2017; Wang et al., 2015). For example, Glasser et al showed that the subject-specific spatial topology of area 55b in relation to the frontal and premotor eye fields substantially diverged from the group average in 11% of subjects (Glasser et al., 2016). In addition, the size of all cortical areas, including large ones like V1, varies by twofold or more across individuals (Amunts, Malikovic, Mohlberg, Schormann, & Zilles, 2000; Glasser et al., 2016). This extensive presence of spatial variability across individuals highlights the need for analysis methods that are adaptive and better able to accurately capture functional regions in individual subjects. One approach that aims to achieve a more accurate subject-specific description of this spatial variability is PROFUMO, which simultaneously estimates subject and group probabilistic functional mode (PFM) maps and network matrices (instead of separate parcellation and mapping steps) (Harrison et al., 2015). In the present study, we show that the spatial variability across subjects captured in these PFMs is strongly associated with behaviour.

Conceptually, network edges are commonly thought of as reflecting coupling strength between spatially separated neuronal populations. However, as discussed above, edge estimates are highly sensitive to spatial misalignments across individuals. Additionally, correlation-based edge estimates are influenced by the amplitudes of localised spontaneous rfMRI fluctuations (Duff, Makin, Smith, & Woolrich, 2017), which have been shown to capture trait variability across subjects, and state variability within an individual over time (Bijsterbosch et al., 2017). These results demonstrate the sensitivity of edge-strength estimates to a wide range of different types of subject variability, and highlight the need to identify which aspects of FC tap directly into behaviourally-relevant population-level variability. Here, we investigate the complex relationships between different features of an rfMRI dataset and also the associations with variability across individuals in terms of their performance on behavioural tests, their lifestyle choices, and demographic information. Using data from the Human Connectome Project (HCP), we provide evidence for systematic differences in the spatial organisation of functional regions. We then use simulations that manipulate aspects of the data to argue that these differences reflect meaningful cross-subject information and drive edge estimates for several common FC approaches.

## Results

### Cross-subject information in fMRI-derived measures

To determine whether a given rfMRI-derived FC measure contains meaningful cross-subject information rather than random variability, we adopted an approach that makes use of the extensive set of behavioural, demographic, and lifestyle data acquired in the HCP. Our first analysis aims to determine which measures obtained from rfMRI and task data most strongly relate to interesting behavioural variability across individuals. Using Canonical Correlation Analysis (CCA), we extracted population modes of cross-subject covariation that represent maximum correlations between combinations of variables in the subject behavioural measures and in the fMRI-derived measures, uncovering multivariate relationships between brain and behaviour. For example, previous work has used CCA on HCP data to identify a mode of population covariation that linked a positive-negative axis of behavioural variables to patterns of FC edge strength (Smith et al., 2015). A specific pattern of connectivity, primarily between “task-negative” (default mode) regions (Raichle et al., 2001), was found to be linked to scores on positive factors such as life satisfaction and intelligence, and inversely associated with scores on negative factors such as drug use. We calculated a CCA score for each subject to represent their position along the population continuum for the latent CCA variable(s), and these scores were subsequently used to visualise variation at both the population extremes (see Figure 2 below), and across the full population continuum (supplementary movies).

We applied a separate CCA analysis for each of the various fMRI-derived measures. The results (Figure 1 and Tables S1 and S2) reveal that highly similar associations with behaviour and life factors occur across a wide range of different fMRI-derived measures. In particular, the results show that spatial features such as PFM subject spatial maps and subject task contrast maps are strongly associated with behaviour. Comparable spatial effects are found when looking at subject-specific parcels in a multimodal parcellation (HCP_MMP1.0; Figure S1) (Glasser et al., 2016). These findings suggest that a large proportion of behavioural information may be represented as cross-subject spatial variability in the functional topography of brain regions.

**Figure 1:**
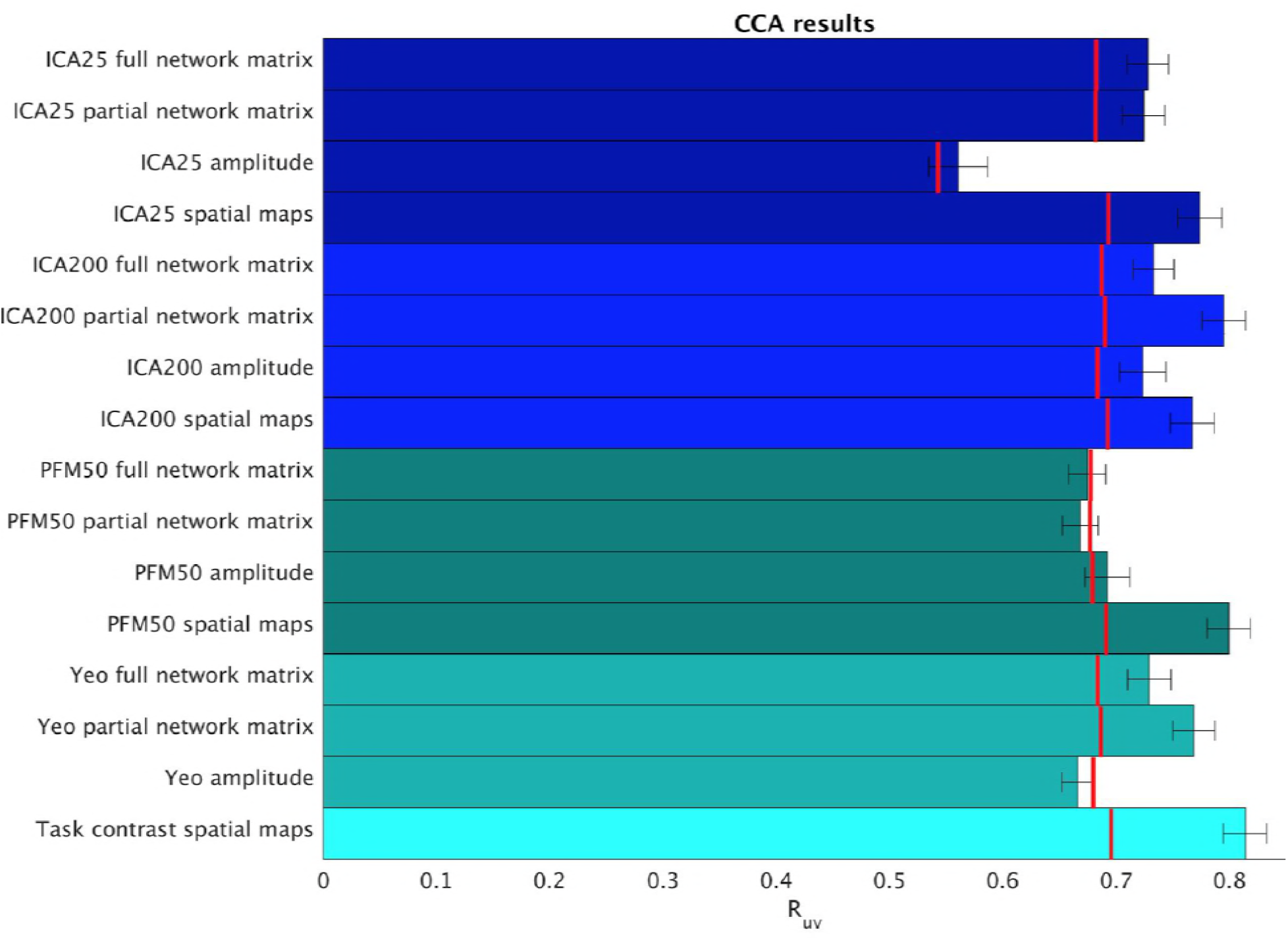
Highly similar associations between behaviour and the brain occur across 16 distinct measures derived from fMRI. This includes network matrices, spatial maps and amplitudes (node timeseries standard deviation) derived from several distinct group-average spatial parcellations/decompositions: ICA decompositions at two scales of detail (dimensionalities of 25 and 200, with “ICA200 partial network matrix” corresponding to the measures used previously(Smith et al., 2015)); a PROFUMO decomposition (PFM; dimensionality 50); an atlas-based hard parcellation (108 parcels(Yeo et al., 2011)), and task contrast spatial maps (86 contrasts). Each bar reports a separate CCA analysis, performed against behaviour/life-factors. A similar mode of variation is found across most of the parcellation methods and different fMRI measures. R_UV_ is the strength of the canonical correlation between imaging and non-imaging measures. Error bars indicate confidence intervals (2.5-97.5%) estimated using surrogate data, and red lines reflect the p<0.002 significant threshold compared with a null distribution obtained with permutation testing (i.e. family-wise-error corrected across all CCA components and Bonferroni corrected across a total of 25 CCAs performed, see Tables S1 and S2 for the full set of results). CCA estimates the highest possible R_uv_ given the dataset; therefore the null distribution for low-dimensional brain data (e.g. ICA 25 amplitude) is expected to be lower than for high-dimensional brain data.

For correlation-based parcellated FC estimates (network edges), a common assumption is that functional coupling is primarily reflected in the edges. However, true network coupling information can in theory be manifested anywhere along a continuum of appearing purely in spatial maps at one extreme (as is the case when performing temporal ICA, where the temporal correlation matrix between components is by definition the identity matrix (Smith et al., 2012)), or purely in edge estimates at the other extreme (as is often assumed to be the case when using an individualized hard parcellation, and would truly be the case if the subjects were all perfectly functionally aligned to the parcellation and contained no useful information in the node timeseries amplitudes). It is likely that the dimensionality of the decomposition may influence this; for example, for a low-dimensional decomposition (into a small number of large-scale networks), much cross-subject variation in functional coupling is likely to occur between sub-nodes of the networks, which is therefore more likely to be represented in the spatial maps, whereas in a higher dimensionality decomposition this information is more likely to be represented in the network matrix. However, the results in Figure 1 show that this CCA mode of population covariation is significantly present in both spatial maps and network matrices for both low and high dimensional decompositions (ICA 25 and 200). Therefore, the potential role of dimensionality is not sufficient to explain the common information present in spatial maps, timeseries amplitudes, and network matrices.

The presence of this behaviourally meaningful spatial variability is somewhat surprising, because these data were aligned using a Multimodal Surface Matching (MSM) approach (Robinson et al., 2014, 2017), driven by both structural and functional cortical features (including myelin maps and resting state network maps). MSM has been shown to achieve very good functional alignment compared with other methods, and particularly compared with volumetric alignment approaches or surface-based approaches that use cortical folding patterns rather than areal features (Coalson, Van Essen, & Glasser, n.d.). However, residual cross-subject spatial variability is still present in the HCP data after the registration to a common surface atlas space (in part due to the constrained parameterisation of MSM and in part due to the underestimation of dual regression subject maps used to drive MSM). In line with this, approaches which are expected to better identify residual subject spatial variability (such as PFM spatial maps and subject task contrast maps) show strong correspondence between spatial variability and behaviour/life-factor measures.

To better understand what spatial features that represent behaviourally-relevant cross-subject information, we visually explored what aspects of the PFM spatial maps contributed to the CCA result in Figure 1 by calculating representative maps at extremes of the CCA mode of population covariation (based on CCA subject scores). The results reveal complex changes in spatial topography (Figure 2 and supplementary movies). For example, comparing left versus right panels shows the right inferior parietal node of the DMN extending farther into the intraparietal sulcus (in the vicinity of area IP1 (Choi et al., 2006; Glasser et al., 2016)) in subjects who score higher on the behavioural positive-negative mode of covariation.

**Figure 2:**
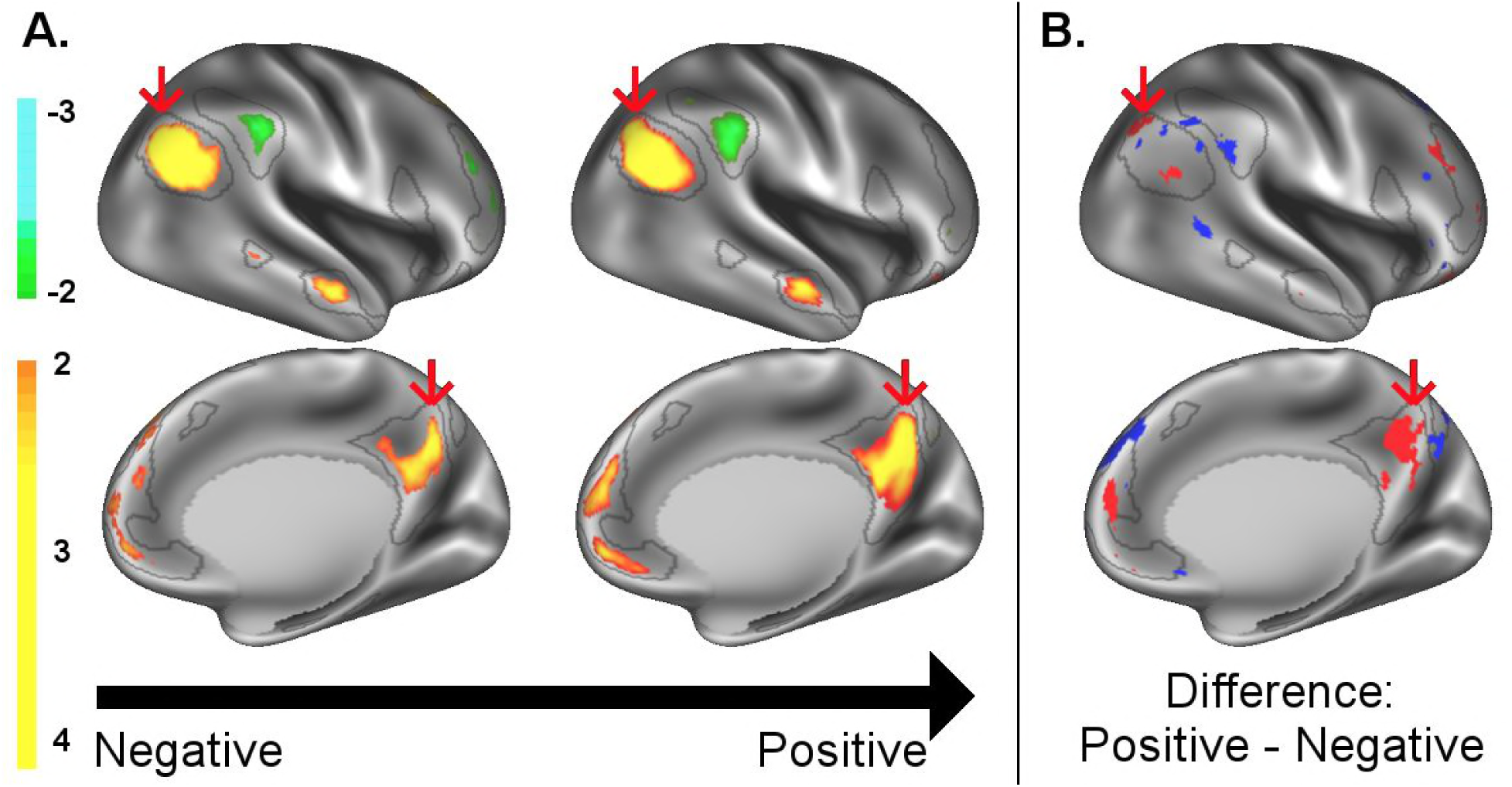
A: representative maps of the two extreme ends (identified based on the low and high extremes along a linearly spaced vector that spans the full range of subject CCA scores) of the CCA mode of population covariation continuum are shown for the default mode network (DMN, the PFM mode that contributed most strongly to the CCA mode of population covariation). The top row shows that the inferior parietal node of the DMN differs in shape and extends into the intraparietal sulcus in subjects who score high on the positive-negative CCA mode (left), compared with subjects who score lower (right). The bottom row shows that medial prefrontal and posterior cingulate/ precuneus regions of the DMN change in size and shape as a function of the CCA positive-negative mode. B: difference maps (positive - negative; thresholded at ± 1) are shown to aid comparison. The representative maps at both extremes are thresholded at ± 2 (arbitrary units specific to the PFM algorithm) for visualisation purposes (the differences are not affected by the thresholding; for unthresholded movie-versions of these maps, please see the supplementary movies which can be downloaded here to aid the review process: https://drive.google.com/drive/folders/0B6J0Q9KXPsNYWmlhTENpa3BKRmc?usp=sharing). The grey contours are identical on the left and right to aid visual comparison, and are based on the group-average maps (thresholded at 0.75). Spatial changes of all PFM modes can be seen in the supplementary movies.

### Spatiotemporal simulations demonstrating potential sources of variability in edges

Figure 1 showed that functionally-relevant cross-subject variability is represented in a variety of different measures derived from both resting state and task fMRI. These widespread similarities in correlations with behaviour across a range of measures invite the question of whether the same type of trait variability is meaningfully and interpretably reflected in a wide range of rfMRI measures, or whether (for example) estimates of network matrices may instead primarily reflect trait variability in spatial topography or amplitude (and not coupling strength). Therefore, we wanted to determine to what extent correlation-based FC measures derived from rfMRI can be influenced by specific aspects of the rfMRI data such as true topography and true coupling. To this end, we created simulated datasets based on the original PFM subjects and/or group spatial maps and timeseries. However, by holding either the individual (simulated) subjects’ spatial maps or the network matrices fixed to the group average we eliminated specific forms of underlying subject variability from the simulated data (Figure 3). We used PFMs in order to generate simulated data because the PROFUMO model separately estimates spatial maps, network matrices and amplitudes, thereby allowing each aspect to be fixed to the group average prior to generating simulated data using the outer product. The aim of the simulation analyses was to determine which features in the rfMRI data are likely to be most strongly reflected in network matrices estimated from rfMRI data. We assess this in terms of the amount of variability across subjects that can be explained, as this is the most relevant application in biomarker studies and in neuroimaging research more generally.

Timeseries were extracted from both the simulated and original datasets, and network matrices were estimated. Each simulated dataset was assessed using three metrics: i) comparing subject-specific simulated and original network matrices *(Z*_network matrix_ in Table 1), ii) comparing cross-subject variability in the simulated and original network matrices (R_correlation_ in Table 1), and iii) determining how much of the cross-subject variability in simulated and original network matrices is behaviourally informative using CCA (see Table 1 legend).

**Figure 3:**
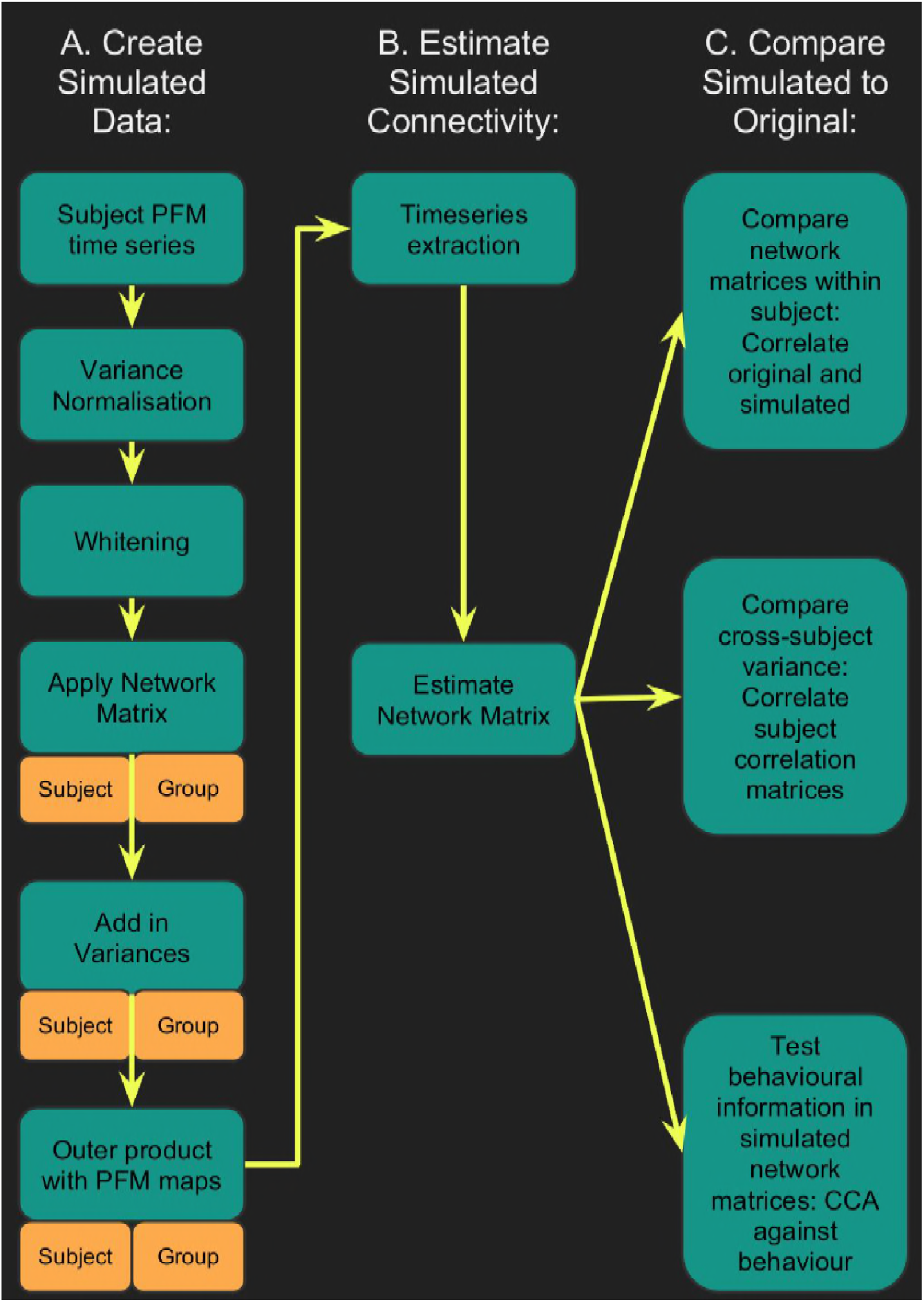
Simulated data is generated for each subject by setting one or more aspects (from the network matrices, node amplitudes and spatial maps) to the group average. Timeseries extraction is performed (using either dual regression against original group ICA maps, or masking against a binary parcellation); network matrices are calculated and compared against network matrices estimated from the original data.

The results (Tables 1, S3, and S4) show that, when the subject-varying aspects of the simulations were exclusively driven by spatial changes across subjects (with the predefined network matrix and amplitudes being identical for all subjects), up to 62% (i.e. square of R_correlation_=0.79 from Table S4 “maps only”) of the cross-subject variance present in the network matrices obtained from the original data was regenerated. Hence, this finding reveals that very similar network matrices can be obtained for any individual subject even if the only aspect of the rfMRI that is varying across subjects is the topographic information in PFM spatial maps. In addition, the variance that can be explained by spatial maps is behaviourally relevant; the CCA results were similarly strong (typically having the same permutation-based p-values) from simulated network matrices driven purely by spatial changes, compared with those obtained from the original dataset.

The influence of amplitudes on FC estimates was relatively minor (less than 2.5% of variance was explained by amplitude in all our simulations; i.e. square of R_correlation_=0.15 from Table 1 “amplitudes only”), although, when amplitudes were combined with spatial maps feeding into the simulations, the amplitudes did in most cases result in an increase in original network matrix regeneration.

**Table 1:** Results from simulated datasets in which one or more of the network matrices, amplitudes and spatial maps are fixed to the group average to remove any subject variability associated with it. Results in each row were driven by variables in which subject variability was preserved, as indicated with ✓ (variables with ‘-’ were fixed to the group average). Results are shown for within-subject correlations between simulated and original z-transformed network matrices (Z_network matrix_), similarities of cross-subject variability represented in simulated and original network matrices (R_correlation_), and for results obtained from the CCA against behaviour (where r_U-V_ is the strength of the canonical correlation between imaging and non-imaging measures, P_U-V_ is the associated (family-wise error corrected) p-value estimated using permutation testing, taking into account family structure, and r_U-Uica_ is the correlation of a CCA mode (subject weights) with the positive-negative mode of population covariation obtained from ICA200 partial network matrices as used in (Smith et al., 2015). For brevity, this Table presents results from full correlation network matrices obtained from a dual regression of ICA 200 maps onto the simulated data (because this approach closely matches previously published findings (Smith et al., 2015)), results for other parcellations are in Table S3 and for partial correlation network matrices in Table S4.

|  | **Simulation driven by true subject variability in:** | **Network matrix** | **Amplitudes** | **Spatial maps** | **Z network matrix** | **R correlation** | **CCA $r_{U-V}$** | **CCA $P_{U-V}$** | **CCA $r_{U-Uica}$** |
| --- | --- | --- | --- | --- | --- | --- | --- | --- | --- |
| ICA | Nothing | - | - | - | -0.0003 | 0.03 | 0.65 | 0.32017 | 0.11 |
| D = 200 | Amps & maps | - | ✓ | ✓ | 1.14 | 0.60 | 0.71 | 0.00001 | 0.52 |
| N=819 | Connectivity only | ✓ | - | - | 0.47 | 0.65 | 0.69 | 0.00028 | 0.40 |
|  | Amplitudes only | - | ✓ | - | 0.22 | 0.15 | 0.69 | 0.00052 | 0.45 |
|  | **Maps only** | - | - | ✓ | **0.78** | **0.54** | **0.72** | **0.00001** | **0.62** |

Given the complex information present in PFM spatial maps, the effect of spatial information on network matrices can result from cross-subject variability in: i) network size, ii) relative strength of regions within a given network, or iii) size and spatial location of functional regions. We performed two further tests to distinguish these influences by thresholding and binarizing the subject-specific spatial maps used to create the simulated data. Maps were either thresholded using a fixed threshold (removing the influence of relative strength), or (separately) using a percentile threshold (removing the influence of relative strength and size, as the total number of grayordinates in binarised PFM maps is fixed across subjects and PFMs). The role of subject-varying spatial maps in driving the resulting estimated network matrices remains strong when highly simplified binarized maps are used to drive the simulations (Table S5), further supporting our interpretation that the results are largely driven by the shape of the functional regions (i.e., variability in the location and shape of functional regions across subjects), rather than by size or local strength.

### Unique contribution of topography versus coupling

The results presented above show that a large proportion of the variance in estimated network matrices is also represented in spatial topography. This suggests either that cross-subject information is represented in both the coupling strength between neural populations and in the ‘true’ underlying spatial topography, or that edge estimates obtained from rfMRI data primarily reflect cross-subject spatial variability (which indirectly drives edge estimates through the influence of spatial misalignment on timeseries extraction, particularly when group parcellations are mapped onto individual subjects in the case of imperfect alignment). To test these hypotheses further, we investigated the unique information contained in spatial maps and network matrices using a set of 15 ICA basis maps derived from HCP task contrast maps (Figure 4A). These basis maps can be thought of as the spatial building blocks that can be linearly combined to create activation patterns for any specific HCP task contrast, and can be considered here to be another functional parcellation.

The advantage of using basis maps derived from task data is that the tasks essentially act as functional localisers that allow for the precise localisation of task-related functional regions within an individual; results at a single-subject level are not influenced in any way, including spatially, by the group results, as they are derived via a temporal task-paradigm analysis, and not via group-level maps. Hence, subject-based task basis maps are the most accurate description of subject-specific locations of functional regions, at least with respect to those regions identifiable from the range of tasks used. Either group-based task basis maps or subject-based task basis maps were entered into a dual regression analysis against subjects’ resting-state fMRI data to obtain network matrices (from dual regression stage 1 timeseries) and rfMRI-based spatial maps (from dual regression stage 2) for each subject (Figure 4B). Subsequently, CCA was performed to determine how well each of the group-based and subject-task-based rfMRI maps and network matrices was able to predict behavioural variability. Furthermore, a ‘partial CCA’ was performed to characterise the unique variance that task rfMRI maps carry over and above network matrices, and vice versa.

The results from the CCAs against behavioural measures show that task rfMRI spatial maps (both subject- and group-based) capture more behavioural information than network matrices (and continue to reach significance in the partial CCA), consistent with the PFM spatial results presented in Figure 1. The strongest partial CCA result was obtained from subject-task-based rfMRI maps (far right in Figure 4C), which are the maps that are expected to contain the most accurate representation of subject-specific functional regions. The results for these spatial maps show the smallest difference between the full and partial CCA results (particularly compared with the spatial maps obtained from the group-task-based rfMRI maps). This suggests that subject variability is more uniquely represented in the spatial information, rather than filtering through into the network matrices. Importantly, this interpretation is supported by the fact that subject-task-based rfMRI network matrices explain the behavioural data considerably less well than group-based task-rfMRI network matrices (difference: p=0.00052 for full network matrices), confirming that spatial information is a significant factor in estimated network matrices.

Taken together, these results show that, while network matrices obtained from dual regression against group-level maps do contain behaviourally relevant cross-subject information, this can be almost completely explained by variability in spatial topographical features across subjects (to the extent that we can detect it). Hence, dual regression network matrices apparently contain little unique information regarding coupling strength that is not also reflected in spatial topographical organisation. However, it is possible that network matrices obtained using parcellation methods and timeseries extraction approaches that are better able to capture subject-specific spatial variability (such as the HCP_MMP1.0 parcellation) do contain unique cross-subject information; further research is needed to test this possibility.

**Figure 4:**
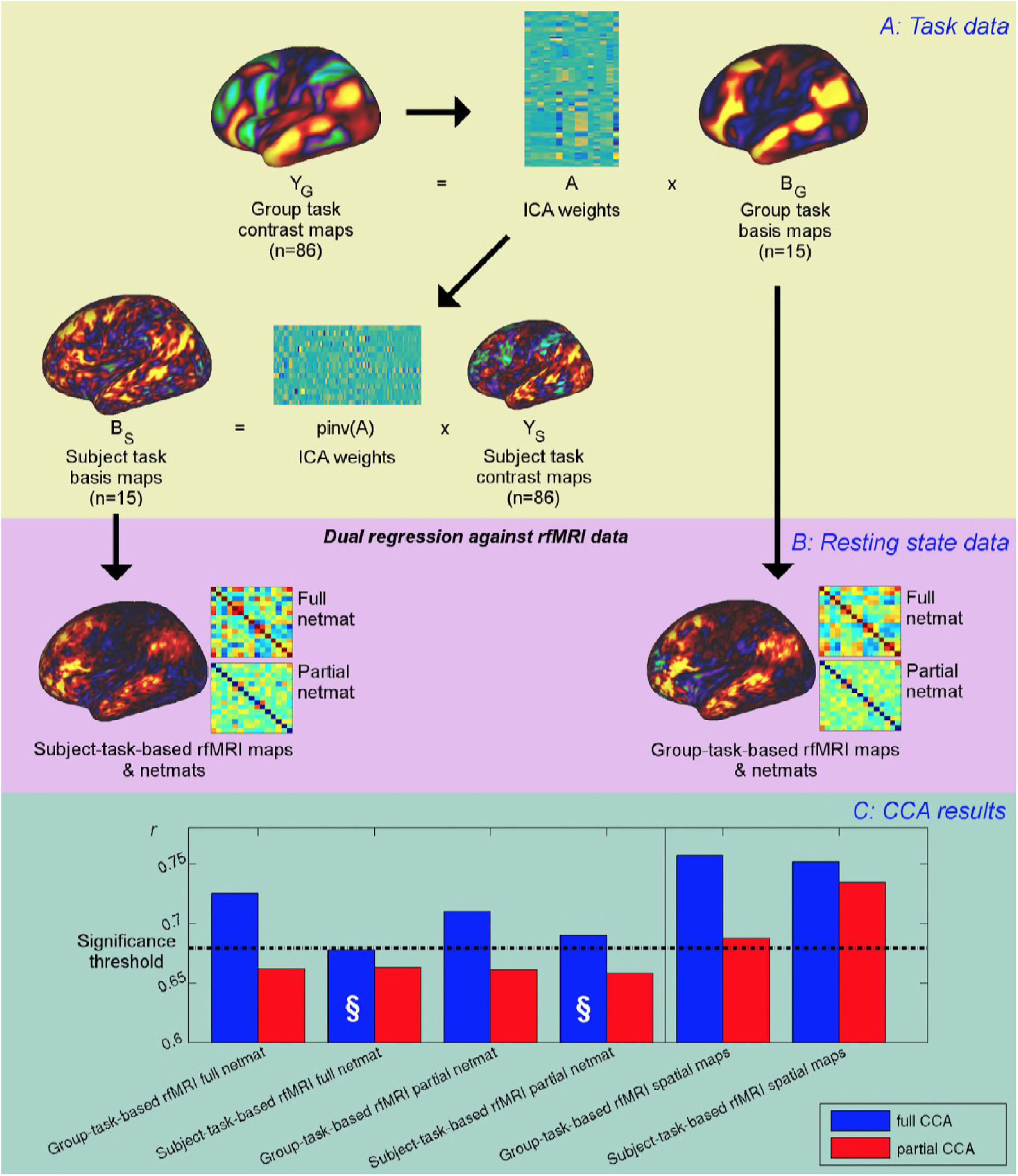
Unique contribution of topography versus coupling. A: Task basis maps are extracted from group-averaged task contrasts using ICA to ensure correspondence of basis maps across subjects. These maps represent the basic building blocks of any activation pattern, and subject task basis maps (obtained by applying the ICA weights to subject task contrast maps) are not influenced by misalignment problems. B: Dual regression against rfMRI data is performed using either the (potentially misaligned) group task basis maps or the (functionally localised) subject task basis maps. C: CCA results of group-task-based rfMRI maps and network matrices and of subject-task-based rfMRI maps and netmats. The results show r_UV_ (i.e., the correlation between the first U and V obtained from the CCA analysis describing the strength of association between the rfMRI and behavioural measures). The null line (i.e., p=0.05 based on permutation testing) is shown as a dotted line at 0.68; results below this line do not reach significance. The blue bars show the main CCA results using the complete data, and the red bars show partial CCA results computed after regressing out any variance that can be explained by network matrices from the spatial maps and vice versa prior to running the CCA. The results show a general decrease in r_UV_ for all measures when comparing partial to full CCA results. The strongest partial CCA result (red bar on right) is found for maps obtained from subject-task-based rfMRI maps, and the associated netmats showed the weakest results (“§”). However, the partial CCA results for the spatial maps (i.e., the red bars on the right) still reach significance. All of the partial CCAs also showed lower r_U-Uica_ compared to the full CCAs (not shown here).

## Discussion

Here, we have identified a key aspect of rfMRI data that directly reflects interesting variability in behaviour and lifestyle across individuals. Our results indicate that spatial variation in the topography of functional regions across individuals is strongly associated with behaviour (Figure 1). In addition, network matrices (as estimated with masking or dual regression against group-level parcellations) reflect little or no unique cross-subject information that is not also captured by spatial topographical variability (Figure 4). This unexpected finding implies that the common interpretation of FC as representing cross-subject (trait) variability in the coupling strength of interactions between neural populations may not be a valid inference (although within-subject state-dependent changes in coupling may still be reflected in FC measures). Specifically, we show that up to 62% of the variance in rfMRI-derived network matrices (a measure commonly taken as a proxy for coupling) can be explained purely by spatial variability. These findings have important implications for the interpretation of FC, and may contribute to a deeper mechanistic understanding of the role of intrinsic FC in cognition and disease (Mill, Ito, & Cole, 2017).

Our findings are consistent with previous research that has highlighted the presence of structured cross-subject spatial variance in both functional and anatomical networks (Glasser et al., 2016; Gordon, Laumann, Adeyemo, Gilmore, et al., 2016; Noble et al., 2015; Sabuncu et al., 2016; Tong, Aganj, Ge, Polimeni, & Fischl, 2017; Xu et al., 2016). Furthermore, recent work has shown that resting state spatial maps can be used to predict task activation maps from individual subjects very accurately (Tavor et al., 2016). Therefore, the presence of behaviourally relevant cross-subject variance in maps of functional (co-)activation in itself is not surprising. However, the fact that the these variations in spatial topographical features capture a more direct and unique representation of subject variability than temporal correlations between regions defined by group parcellation approaches (coupling), was unexpected. The implication of this finding is that the cross-subject information represented in commonly adopted ‘connectivity fingerprints’ largely reflects spatial variability in the location of functional regions across individuals, rather than variability in coupling strength (at least for methods that directly map group-level parcellations onto individual data). Specifically, our partial CCA results (Figure 4) show that network matrices (as often estimated) contain little unique trait-level cross-subject information that is not also reflected in the spatial topographical organisation of functional regions.

How the functional organisation of the brain is conceptualised and operationally defined is of direct relevance to the interpretation of these findings. Some hard parcellation models of the human cortex (such as the Gordon and Yeo parcellations (Gordon, Laumann, Adeyemo, Huckins, et al., 2016; Yeo et al., 2011)) aim to fully represent connectivity information in the edges (i.e. correlations between node timeseries). Thus, hard parcellations of this type assume piecewise constant connectivity within any one parcel (i.e. each parcel is assumed to be homogeneous in function, with no state- or trait-dependent within-parcel variability in functional organisation). In contrast, the HCP_MMP1.0 multimodal parcellation presumes within-area uniformity of one or more major features, but overtly recognizes within-area heterogeneity in other features, including connectivity, most notably for distinct body part representations (‘sub-areas’) of the somatomotor complex. Soft parcellation models (such as PROFUMO (Harrison et al., 2015)) allow for the presence of multiple modes of (potentially overlapping) functional organisation. Therefore PFMs represent connectivity information through complex interactions between amplitude and shape in the spatial maps, and through network matrices. Our findings show that both the PROFUMO and the multimodal parcellation models successfully capture behaviourally-relevant cross-subject spatial variability (Table S2), but that the precise location of where this spatial variability is represented overlaps only modestly between the two approaches (Figure S1). Given the differences in the key assumptions made by the two models (i.e. binary parcellation versus multiple modes of functional organisation), this is not unexpected. However, it does highlight the need for further research into the optimal representation of (subject-specific) functional organization in the brain.

For most of the results presented in this work, we estimated spatial information using functional data (either resting or task fMRI data). While a comprehensive investigation of related anatomical features is beyond the scope of this work, we did identify significant correlations between fractional surface area size and subject CCA weights (Figure S1). This result suggests that anatomical variability in the cortical extent of a number of higher level sensory and cognitive brain regions may contribute to the overall findings presented here. Further research into the relationship between structural features and functional connectivity measures, and their contribution to trait-level subject variability is needed to test this hypothesis.

The results presented are of relevance to a large variety of approaches used to study connectivity. For example, our simulation results (Tables 1, S3, and S4) reveal similar results regardless of whether we adopt a dual-regression or a masking approach to obtain timeseries, and the findings also do not differ qualitatively according to whether full or partial correlation is used to estimate network matrices. Therefore, our findings are relevant to any approach that is based on timeseries extracted from functional regions defined at the group-level (including graph theory methods and spectral analyses). The implications of this work may also extend beyond resting-state fMRI. For example, generative models such as dynamic causal modelling (DCM) are increasingly used to stratify patient populations (Brodersen et al., 2014), and to achieve predictions for individual patients (Stephan et al., 2017). Previous work has shown that including parameters for the position and shape of functional regions in individual subjects into the model improves DCM results and better differentiates between competing models (Woolrich, Behrens, & Jbabdi, 2009). At the moment, it is unknown to what extent cross-subject variability observed with these timeseries-based fMRI metrics reflects true coupling between neural populations, rather than being indirectly driven by spatial variability and misalignment. Going forward, it is important to disambiguate the influence of spatial topography, to enable the estimation of fMRI measures that uniquely reflect coupling strength between neural populations.

Significant advances have already been made in recent years in order to tackle the issue of spatial misalignment across individuals. For example, the HCP data used in this work were spatially aligned using the multimodal surface mapping (MSM) technique, which achieves very good functional alignment by using features that are more closely tied to cortical areas (although note that, since the time of the HCP release, refinements to the [regularisation of the] MSM algorithm have resulted in further improvements in the observed functional alignment of HCP data (Robinson et al., 2014, 2017)). Therefore, gross misalignment is unlikely to play a role in our results. In fact, some of the behaviourally relevant variability may have been ‘corrected’ in the MSM pipeline prior to our analyses (indeed, the same positive-negative mode of population covariation is identified when running the CCA on MSM warp fields; Table S1; and the fractional surface area results in Table S2 and Figure S1A reflect the full variability from native space, and is therefore not affected by the alignment accuracy). Therefore, it is possible that the degree to which spatial information may influence FC estimates varies considerably across studies, depending on the spatial alignment algorithm that was used, and the amount of subject spatial variability this has removed. It is encouraging that significant efforts have recently gone into the methods for more accurately estimating the spatial location of functional parcels in individual subjects in recent years (Chong et al., 2017; Glasser et al., 2016; Gordon, Laumann, Adeyemo, Huckins, et al., 2016; Hacker et al., 2013; Harrison et al., 2015; Varoquaux, Gramfort, Pedregosa, Michel, & Thirion, 2011; Wang et al., 2015). The present results highlight the importance of such advances, and call for the continued development, comparison, and validation of such approaches.

In conclusion, we have demonstrated that spatial topography of functional regions are strongly predictive of variation in behaviour and lifestyle factors across individuals, and that timeseries-based methods (as often estimated based on group-level parcellations) contain little unique trait-level information that is not also explained by spatial variability.

## Materials and Methods

### Dataset

For this study we used data from the Human Connectome Project S900 release (820 subjects with fully complete resting-state fMRI data, 452 male, mean age 28.8 ± 3.7 years old) (Van Essen et al., 2013). Data were acquired across four runs using multiband echo-planar imaging (MB factor 8, TR = 0.72 sec, 2mm isotropic voxels) (Moeller et al., 2010; Ugurbil et al., 2013). Data were preprocessed according to the previously published pipeline that includes tools from FSL, Freesurfer, HCP’s Connectome Workbench, multimodal spatial alignment driven by myelin maps, resting state network maps, and resting state visuotopic maps (“MSMAll”), resulting in data in the grayordinate coordinate system (Fischl, Sereno, & Dale, 1999; Glasser et al., 2013, 2016; Jenkinson, Beckmann, Behrens, Woolrich, & Smith, 2012; Marcus et al., 2013; Robinson et al., 2014; Smith, Beckmann, et al., 2013). ICA-FIX-cleanup was performed on individual runs to reduce structured noise (Griffanti et al., 2014; Salimi-Khorshidi et al., 2014).

### Data Availability

HCP data are freely available from https://db.humanconnectome.org. The version of MSMAll that is compatible with the approach implemented for the alignment of HCP data can be found here: http://www.doc.ic.ac.uk/~ecr05/MSM_HOCR_v2/ (Robinson et al., 2017). Matlab code used in this work can be found here: https://github.com/JanineBijsterbosch/Spatial_netmat.

### Inferring functional modes

In order to obtain estimates of the spatial shape and size of functional networks for every subject, we decompose the HCP data into a set of probabilistic functional modes (PFMs) via the PROFUMO algorithm (Harrison et al., 2015). A set of *M* PFMs describe each subject’s data (*G* grayordinates; *T* time points; *D*_*s*_∈*R*^*V×T*^) in terms of a set of subject-specific spatial maps (*P* _*s*_∈*R*^*V×M*^), amplitudes (*h*_*s*_∈*R*^*M*^) and timecourses (*A*_*s*_∈*R*^*M×T*^), all of which are linked via the outer product model:

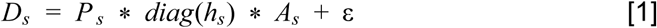

These subject-specific decompositions are linked by a set of hierarchical priors. In the spatial domain, the group-level parameters encode the grayordinate-wise means, variances and sparsity of the subject maps, while in the temporal domain, the group-level priors constrain the subject-level network matrices (note that the component amplitudes and hierarchical priors are recent extensions to the PFMs model and were not included in the original PROFUMO paper (Harrison et al., 2015)). The PROFUMO framework gives us sensitive estimates of key subject-level parameters, while ensuring that there is direct correspondence between PFMs across subjects.

PROFUMO was run on the rfMRI data from all 820 subjects with a dimensionality of 50 PFMs. Importantly, the signal-subspace of any given subject’s dataset can be straightforwardly reconstructed from a set of modes via equation [1], and this can be used to generate the simulated data as described below.

### Canonical Correlation Analysis (CCA)

For the ICA decompositions, amplitudes were estimated for each subject and component as the temporal standard deviation of the timeseries obtained from stage 1 of a dual regression analysis. Full and regularised partial correlation matrices were also calculated from these timeseries. The Tikhonov regularisation rho used during estimation of the partial correlation matrices was set to 0.01 for the ICA 25, 200 and PFM data (according to previous optimisation results). For high dimensional parcellations (Yeo and HCP_MMP1.0), the rho was optimised by finding the maximum correlation between subject and group-average (using rhoe = 0.01) network matrices across a range of rho (0.01:0.5), leading to rho=0.03 for Yeo and rho=0.23 for HCP_MMP1.0 results. Lastly, the subject spatial maps obtained from stage 2 of a dual regression analysis were used. Similarly, for the PROFUMO decomposition, the PFM amplitudes, subject spatial maps and timeseries were used. For the HCP_MMP1.0 spatial results, either group-level or subject-specific node parcellations were used(Hacker et al., 2013). The subject-specific parcellations contain missing nodes (parcels) in some subjects (Glasser et al., 2016). Hence, for partial network matrices, the rows and columns in the covariance matrix were set to the scaled group average prior to inverting the covariance matrix. In the resulting network matrices, the rows and columns relating to missing nodes were set to the group average (for both partial and full network matrices). Before performing CCA, missing nodes were accounted for by estimating the subject-by-subject covariance matrix one element at a time, ignoring any missing nodes for any pair of subjects. The nearest valid positive-definite covariance matrix was subsequently obtained using nearestSPD in Matlab (http://uk.mathworks.com/matlabcentral/fileexchange/42885-nearestspd), prior to performing singular value decomposition as described below.

Each CCA analysis finds a linear combination of behavioural and life-factor measures (V) that is maximally correlated with a linear combination of rfMRI-derived measures (U) (Hotelling, 1936): *Y* * *A* = *U ~ X* * *B* = *V*. Y is the set behavioural measures, and X are the rfMRI-derived measures (i.e. spatial maps, or network matrices, or signal amplitudes). A and B are optimised such that the correlation between U and V is maximal. Summary measures from CCA include the correlation between (paired columns of) U and V, and the associated p-values (derived from permutation testing over n=100,000 permutations) for the first one or more CCA modes.

To create the inputs to the CCA, a set of nuisance variables were regressed out of both the behavioural measures and the amplitudes, network matrices and spatial maps, as done in(Smith et al., 2015). Subject covariance matrices were subsequently estimated for the amplitudes, network matrices and for all spatial maps (by summing the covariance matrices of individual spatial maps). Then a singular value decomposition was performed on the subject covariance matrices and the first 100 eigenvectors were entered into the CCA (either against 100 eigenvectors obtained from behavioural variables as explained in (Smith et al., 2015), or to compare PFM spatial maps directly to ICA partial correlation matrices).

In addition to reporting the CCA results for the strength of the canonical correlation between imaging and non-imaging measures and the associated p-value (r_U-V_ and P_U-V_), we also report the correlation between the CCA subject weights and the weights for the ICA 200 partial network matrices (r_U-Uica_). The reason for including this correlation is to facilitate direct comparison to previously published CCA results from HCP data (Smith et al., 2015). However, this earlier finding should not be taken as the gold standard CCA result. The r_U-Uica_ correlation we report is the maximum correlation found between the first CCA mode from the ICA 200 partial network matrices, and any of the 100 modes of population covariation obtained for the comparison CCA result (i.e., the maximum correlation may not be with the strongest CCA mode).

Confidence intervals for CCA results in Table 1 were obtained using surrogate data for both the brain-based CCA input matrix and the behaviour CCA input matrix. To generate the surrogate data, row and column wise correlations of the original CCA input matrices were maintained using a multivariate normal random number generator (mvnrnd.m in Matlab). A total of 1000 instances of surrogate data were used to obtain 2.5-97.5% confidence intervals around r_U-V_.

For visualisation and interpretation purposes, we created movies of the spatial variability along the axis of the behavioural CCA mode of population covariation. For this, we took the U resulting from the CCA between PFM spatial maps and behaviour, and created a linearly spaced vector that spans just over the full range of U (extending beyond the lowest and highest measured subject score by 10% of the full range). As the CCA is linear, it is straightforward to project a set of U values back to form a rank-one reconstruction of the original space, which in this case is a set of spatial maps. This sequence of spatial maps is an approximation to the spatial variability that is encoded along the previously reported positive-negative axis. These are used as the frames for supplementary movies 1-9, and for the illustrative examples shown in Figure 1 in the main manuscript (and Figure S2 based on the CCA mode obtained from a CCA performed on PFM spatial maps against ICA partial correlation matrices).

### Creating simulated data

In order to create simulated datasets for each subject, we took the outer product between PFM spatial maps and timeseries. Compared with data that is completely simulated, this approach has the advantage of keeping many features in the data (such as the types of structured noise that are present, the signal-to-noise ratio, and the autocorrelation structure), while still achieving investigator control of specific aspects of interest. Data from each run (1200 time points) was processed separately through the simulation pipeline, including the following steps:

#### Timeseries processing

##### Variance normalisation

Each original PFM subject timecourse was set to unit variance, and the variances were retained.

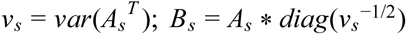

##### Whitening

The ZCA whitening transform (Bell & Sejnowski, 1997) was used to remove any correlations between timeseries:

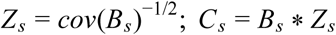

##### Network matrix application

Timeseries were modified such that the induced correlation matched a pre-specified structure.: *D*_*s*_ = *C*_*s*_ * *α*. In the simulations that use a fixed group network matrix, this pre-specified correlation structure was estimated by projecting the S900 group average HCP dense connectome (following Wishart Rolloff) onto the group PFM spatial maps.

##### Restore variances

At this stage the variances of the original timeseries are restored: 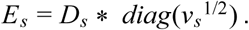. This gives a set of simulated timeseries *E*_*s*_ which have all the same properties as the reference timeseries (*A*_*s*_), except for their correlation structure.

##### Pseudo-PFM generation

We modify the inferred PFMs by selectively setting some of the parameters to their group averages. For example, if we set 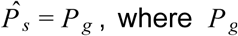, where *P*_*g*_ is the mean over all 820 subject maps, then we can eliminate any spatial variability across subjects. Similarly, we can set the temporal correlations to a fixed group mean using the procedure described above to remove any variability in FC across subjects. In order to remove amplitude variability across subjects, we add in group averaged variances instead of the subject variances. These simulated PFMs are then described by the simulated maps, amplitudes and timeseries, namely 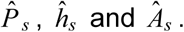

##### Data reconstruction

Finally, the full data can be reconstructed as per [1]: 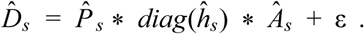. Spatio-temporally white-noise (with variance matched to the original data) is added to the activity described by the simulated modes to give a dataset that preserves the properties of the original data, but, crucially, one where we have direct control over where in the model subject variability can appear.

Once the simulated data is generated for each run, we extracted timeseries from both the simulated and original data using two different approaches that are commonly adopted in the literature. Dual regression analysis was performed using the group ICA maps that were estimated using the (original) HCP group data, and that are freely available with the S900 data release (www.humanconnectome.org). Two dimensionalities were tested, so for each simulated dataset dual regression was performed against 25 and against 200 group ICA components (Figure 5B). The timecourses estimated in stage 1 of the dual regression analysis were used to compute network matrices (Filippini et al., 2009; Nickerson, Smith, Öngür, & Beckmann, 2017). Mean timeseries were also extracted from a set of 109 binary regions of interest (ROIs) based on the Yeo parcellation, and from the HCP_MMP1.0 group parcellations and individual subject parcellations (Glasser et al., 2016). The 109 Yeo ROIs were obtained from the 17-network parcellation(Yeo et al., 2011), by separating each of the 17 networks into individual contiguous regions that had a surface cluster area of at least 20 mm^2^. Timecourses were used to estimate full and regularised partial correlation network matrices using FSLnets (https://fsl.fmrib.ox.ac.uk/fsl/fslwiki/FSLNets). Z-transformation was applied to the network matrices before further comparisons. The network matrices derived from simulated data are compared against network matrices calculated from the original data as described below.

Firstly, we compare the simulated network matrix to the original network matrix for each subject, to determine how similar the measured FC is. For each subject the node-by-node full or regularised partial network matrix estimated from the simulated data is reshaped into a single column after removing the diagonal and is correlated against the reshaped original estimated network matrix. Prior to reshaping the simulated and original network matrices, the respective group average network matrix (simulated or original) is subtracted from the subject network matrix, so that the subsequent correlation is sensitive to the unique subject variability instead of being driven by the group connectivity patterns. As such, a correlation coefficient between demeaned simulated and original network matrices is estimated for each subject. The Fisher r-to-z transform was applied to these correlations before averaging across subjects. This first test assesses how different a subject is from the group (and the similarity of this difference between original and simulated network matrices), and therefore does not test for cross-subject variability.

Secondly, the subject-by-subject correlation matrix was estimated from the subject-wise simulated network matrices. Again, this matrix was reshaped into a vector after discarding the diagonal and was correlated against the reshaped subject-by-subject correlation matrix obtained from the original network matrices. The aim of this test was to directly compare the cross-subject variability present in the simulated and original data, which is very important given that variability across subjects is typically of primary interest in FC research. Hence, this analysis aims to compare the cross-subject variability in original or simulated network matrices, as opposed to comparing the similarity of original and simulated network matrices within an individual subject (as is the case for the preceding approach).

The last test of the simulated network matrices was to perform a CCA against the set of behavioural and life-factor measures (Smith et al., 2015). A CCA was performed on the simulated network matrices against the subject behavioural measures as described below. To asses the CCA results, we report the correlation between U and V (for the first, strongest mode of population covariation), the associated permuted p-value (n=100,000 permutations, respecting family structure), and the maximum correlation between any of the simulated U and the first U obtained when using the original ICA 200 dimensionality partial network matrices describing the positive-negative mode of covariation (Smith et al., 2015).

### Simulations with further spatial map modulations

The PFM subject spatial maps contain a relatively complex set of information. This may include relative differences in amplitude in different brain regions that are part of the same mode, which effectively reflect connectivity rather than spatial shape and size. In order to exclude these potential connectivity-related aspects of the spatial maps and isolate the role of spatial shape, we simplified the spatial maps for some of the simulations presented. For this, the spatial maps were thresholded at a very liberal threshold of 1 (arbitrary units specific to the PFM algorithm) and binarized. The sign was retained such that grayordinates in the subject PFM maps with values >1 were set to 1 and grayordinates with values <-1 were set to −1 and all others to zero. A liberal threshold was purposefully used as we wanted to retain extended (broad, low) shape information, and just remove any information encoded in the (relative) grayordinate amplitudes. Using a fixed threshold across subjects retains cross-subject variability in the size of networks. To further remove this source of information and focus purely on the shape of networks, we applied a percentile threshold such that the size of networks is fixed across subjects (grayordinates > 95th percentile set to 1 and grayordinates < 5th percentile set to −1, leading to each individual PFM map having the same size of 4564 1s and 4564 −1s across all subjects). The results of simulations where the maps were modulated in this way prior to calculating the simulation’s space-time outer product are presented in supplementary Table S5, including results for which the maps were both thresholded and binarized, percentile thresholded and binarized, and also results for maps that were thresholded (at 1) but not binarized.

### Comparing cross-subject similarities between different types of imaging measures

Given that variability between subjects is of primary interest in rfMRI research, this analysis aimed to directly compare the cross-subject variability present in a range of measures obtained from the original data. Between-subject correlation matrices were calculated from network matrices (ICA25, ICA200 and PFM50), from PFM amplitudes and from spatial maps (ICA25 and ICA200 dual regression stage 2 spatial maps, and PFM50 spatial maps). These subject by subject correlation matrices were reshaped after discarding the diagonal, and full and partial correlations were calculated between the subject correlation matrices (Figure S3).

### Unique contribution of topography versus coupling

To obtain a basis set of spatial maps based on task contrast data, we performed a spatial ICA (with a dimensionality of 15) on the concatenated group-averaged task contrast maps (a total of 86 maps, 47 of which are unique). Spatial ICA was performed on the group-average task contrasts maps to avoid the correspondence problem that would arise if ICA were applied separately to individual subject task contrast maps. This resulted in a set of ICA weights (15*86), which describe the contribution of each task contrast map to each extracted ICA component. The outer product of these weights with either the group-averaged contrast maps or the corresponding subject-specific contrast maps was used to obtain maps to drive subsequent dual regression analysis. Dual regression analysis (driven by either group-averaged or subject-specific task basis maps after normalising the maximum of each subject and component map to 1) was run against subject resting state data to obtain timeseries and maps. CCA against behaviour was performed separately on the resulting network matrices and spatial maps as described above. Additionally, partial CCA was performed to determine the unique information contained in network matrices and in spatial maps. For this, any variance explained by network matrices was regressed out of the spatial maps and vice versa (i.e. was ‘partialled out’), before running the “partial CCA”. Specifically the 100 eigenvectors used as the input matrix to the CCA (as explained above and following (Smith et al., 2015)) for partial network matrices were regressed out of the 100 eigenvectors for the spatial maps before running CCA, or conversely the 100 eigenvectors for spatial maps were regressed out of the 100 eigenvectors for the network matrices before running CCA.

## Acknowledgements

Data were provided by the Human Connectome Project, WU-Minn Consortium (Principal Investigators: David Van Essen and Kamil Ugurbil; 1U54MH091657) funded by the 16 NIH Institutes and Centers that support the NIH Blueprint for Neuroscience Research; and by the McDonnell Center for Systems Neuroscience at Washington University. CFB acknowledges support from The Netherlands Organization for Scientific Research (NWO, grant no 864.12.003). We are grateful for funding from the Wellcome Trust (grants 098369/Z/12/Z and 091509/Z/10/Z). The Wellcome Centre for Integrative Neuroimaging is supported by core funding from the Wellcome Trust (203139/Z/16/Z).

## Competing interests

The authors declare no competing financial interests.

## Supplementary Material

### Cross-subject information in fMRI-derived measures

Figure 1 in the main manuscript shows results from separate canonical correlation analyses between the set of lifestyle and behavioural measures and a variety of different fMRI-derived features. Table S1 below shows a more detailed and complete version of Figure 1.

**Table S1:** Highly similar associations between behaviour and the brain can be found across a wide range of different measures derived from fMRI. We included a set of network matrices, spatial maps and amplitudes (node timeseries standard deviation) derived from several distinct group-average spatial parcellations/decompositions: from ICA decompositions at two scales of detail (dimensionalities of 25 and 200); a PROFUMO decomposition (PFM; dimensionality 50); an atlas-based hard parcellation (108 parcels^9^); task contrast spatial maps (86 contrasts); and MSM warp fields from native space to MSMAll aligned data (from estimate_metric_distortion; https://github.com/ecr05/MSM_HOCR_macOSX/blob/master/src/MSM/estimate_metric_distortion.cc). Each row reports a separate CCA analysis, performed against behaviour/life-factors. A very similar mode of variation is found across most of the parcellation methods and different fMRI measures. r_U-V_ is the strength of the canonical correlation between imaging and non-imaging measures (confidence intervals estimated using surrogate data), P_U-V_ is the associated (family-wise error corrected) p-value estimated using permutation testing, taking into account family structure, and r_U-V_ CI is the 2.5-97.5% confidence interval estimated using surrogate data. r_U-Uica_ is the correlation of a CCA mode (subject weights) with the positive-negative mode of population covariation obtained from ICA200 partial network matrices as used in^30^, and is therefore defined to be 1 in the row containing the results from that CCA. The r_U-Uica_ result was included because it shows whether different metrics are associated with similar or distinct behavioural modes of population covariation (one may expect different rfMRI measures to be associated with distinct aspects of behaviour).

| | | $r_{U-V}$ | $r_{U-V}$ CI<br>2.5-97.5% | $P_{U-V}$ | $r_{U-Uica}$ |
| --- | --- | --- | --- | --- | --- |
| **ICA**<br>**d=25**<br>**N=819** | Full correlation network matrix | 0.73 | 0.71-0.75 | 0.00001 | 0.43 |
|  | Partial correlation network matrix | 0.72 | 0.71-0.74 | 0.00001 | 0.42 |
|  | Amplitudes | 0.56 | 0.54-0.59 | 0.00002 | 0.37 |
|  | Spatial maps | 0.77 | 0.76-0.80 | 0.00001 | 0.78 |
| **ICA**<br>**d=200**<br>**N=819** | Full correlation network matrix | 0.73 | 0.72-0.75 | 0.00001 | 0.56 |
| | Partial correlation network matrix | 0.79 | 0.78-0.82 | 0.00001 | $\triangle 1$ |
|  | Amplitudes | 0.72 | 0.70-0.74 | 0.00001 | 0.64 |
|  | Spatial maps | 0.77 | 0.75-0.79 | 0.00001 | 0.78 |
| **PFM**<br>**d=50**<br>**N=819** | Full correlation network matrix | 0.67 | 0.66-0.69 | 0.00482 | 0.31 |
|  | Partial correlation network matrix | 0.67 | 0.65-0.69 | 0.01774 | 0.34 |
|  | Amplitudes | 0.69 | 0.67-0.71 | 0.00006 | 0.29 |
|  | Spatial maps | **0.80** | 0.78-0.82 | 0.00001 | **0.81** |
| **Yeo**<br>**d=108**<br>**N=819** | Full correlation network matrix | 0.73 | 0.71-0.75 | 0.00001 | 0.60 |
|  | Partial correlation network matrix | 0.77 | 0.76-0.79 | 0.00001 | 0.69 |
|  | Amplitudes | 0.67 | 0.65-0.68 | 0.05546 | 0.37 |
| **Task**<br>**N=790** | Contrast spatial maps (d=86) | **0.81** | 0.79-0.83 | 0.00001 | 0.57 |
| **Warp**<br>**N=819** | Warp field from native space to<br>MSMAll alignment | 0.72 | 0.70-0.74 | 0.00001 | 0.53 |

The results in Table S1 reveal that subject-specific PFM spatial maps are strongly associated with cross-subject variability in lifestyle and behaviour. A visual representation of the spatial changes along the continuum of the CCA population mode of covariation for all 32 signal PFMs can be found in the supplementary movies. These movies show changes in spatial topography in representative unthresholded maps. The order of the PFMs reflect how strongly each PFM contributed to the CCA result (i.e., the first movie contains the 4 PFMs that contributed most strongly).

### Comparing PFM and HCP_MMP1.0 spatial results against surface area

Direct comparison between the results in Figure 1 / Table S1 and the HCP_MMP1.0 parcellation (a subject-specific multimodal hard parcellation; 360 parcels^11^) and against associated fractional surface area (in native space as a ratio to total surface area, for each of the 360 parcels in the HCP_MMP1.0 parcellation) is challenging due to the large difference in the number of subjects (n=819 for Table S1 and n=441 for HCP_MMP1.0). Therefore, we have included an analysis on all PFM metrics in a reduced number of subjects (the same n=441 subjects) in order to facilitate direct comparison between these two recent parcellation approaches that both aim to achieve accurate detection of subject-specific spatial boundaries (Table S2). These results show that spatial features from a variety of sources (surface area, multimodal parcellation and PFMs) are strongly associated with measures of behaviour and lifestyle. Also note that network matrices obtained by the HCP_MMP1.0 parcellation are more predictive of behaviour than are PFM network matrices.

**Table S2:**
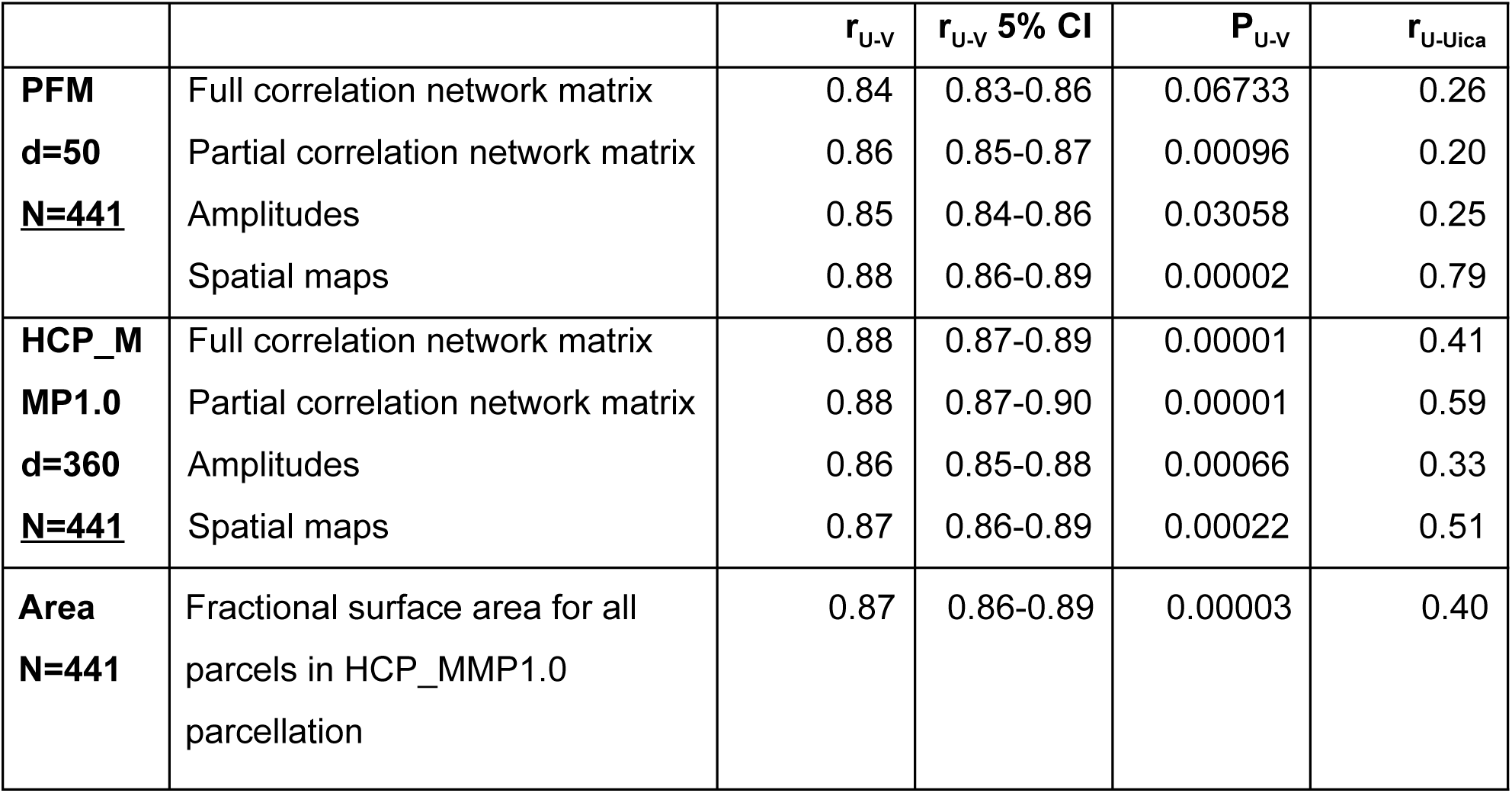
The r_U-V_ results here are inflated in comparison to the results presented in Table S1 (due to increased overfitting as a result of the parcellation only being available in 441 subjects compared with 819 subjects included for the other CCAs), but the associated P_U-V_ can (to some extent) be used for comparison. Therefore, this Table compares PFM (d=50), HCP_MMP1.0 (d=360), and fractional surface area (the fraction of cortex occupied by each area in the multimodal HCP_MMP1.0 parcellation) on the same set of 441 subjects.

The two rfMRI parcellation methods included in Table S2 (HCP_MMP1.0 and PFM) explicitly aim to capture cross-subject variability in the spatial location of functional regions. The subject spatial maps estimated by both methods are strongly associated with cross-subject behavioural variability (when matching the sample size r_U-V_ did not significantly differ, and subject weights of the strongest CCA results were moderately correlated r_U-U_=0.55). Therefore, it is of interest to compare these results in more detail, to determine whether cross-subject variability is represented similarly for the two approaches. Furthermore, given that fractional surface area (the fraction of cortex occupied by each area in the multimodal HCP_MMP1.0 parcellation) was also strongly predictive of behaviour (Table S2), we investigated the potential relationship between rfMRI-based PFM weights, multimodally-defined cortical areal boundaries (HCP_MMP1.0 parcellation), and structural variation in fractional surface area. To this end, we averaged CCA subject weights obtained from two separate CCA results (PFM spatial maps - behaviour, and HCP_MMP1.0 spatial maps - behaviour). These averaged subject weights were subsequently correlated against fractional surface area, and against subject-specific PFM and HCP_MMP1.0 spatial maps (grayordinate-wise), to investigate which brain regions contribute strongly to the association with behaviour, and to compare these localised effects across methods/modalities.

Correlations between fractional area and behaviour were highly consistent between left and right hemispheres, and revealed relatively high correlations in higher order sensory and cognitive regions (Figure S1A). Specifically, bilaterally significant (FDR corrected p<0.05) positive associations between larger surface area and higher scores on the positive-negative mode of population covariation were found in area POS2 of the posterior cingulate cortex and in area IPS1 of the dorsal visual processing stream; bilaterally significant negative correlations were identified in the cingulate motor area 24dv, premotor area 6r, and inferior parietal cortex (areas PFt, PFm, PGi). Qualitative comparison between the spatial localisation of strongest correlations with behaviour across all three datasets reveals that many regions that contribute strongly in either the HCP_MMP1.0 or in the PFM individual subject spatial estimates spatially overlap or adjoin cortical areas in which fractional surface area was also closely linked to behaviour (Figure S1B). This qualitative finding suggests that differences in regional surface area may drive many of the results presented in this work, although further research is needed to confirm this interpretation. The cortical localization of strong associations with behaviour do not closely overlap between PFMs and the HCP_MMP1.0 parcellation (i.e. red and blue regions in Figure S1B and un-thresholded maps in Figure S1C/D). This lack of exact correspondence of the representations of cross-subject variability may reflect differences between the HCP_MMP1.0 and PROFUMO models (the former being a hard parcellation with no overlap between parcels, and the latter being a soft parcellation that includes complex and often overlapping networks), and differences in the data types driving the parcellation (PROFUMO being driven by rfMRI data only, and the HCP_MMP1.0 parcellation being driven by data from multiple different modalities).

**Figure S1:**
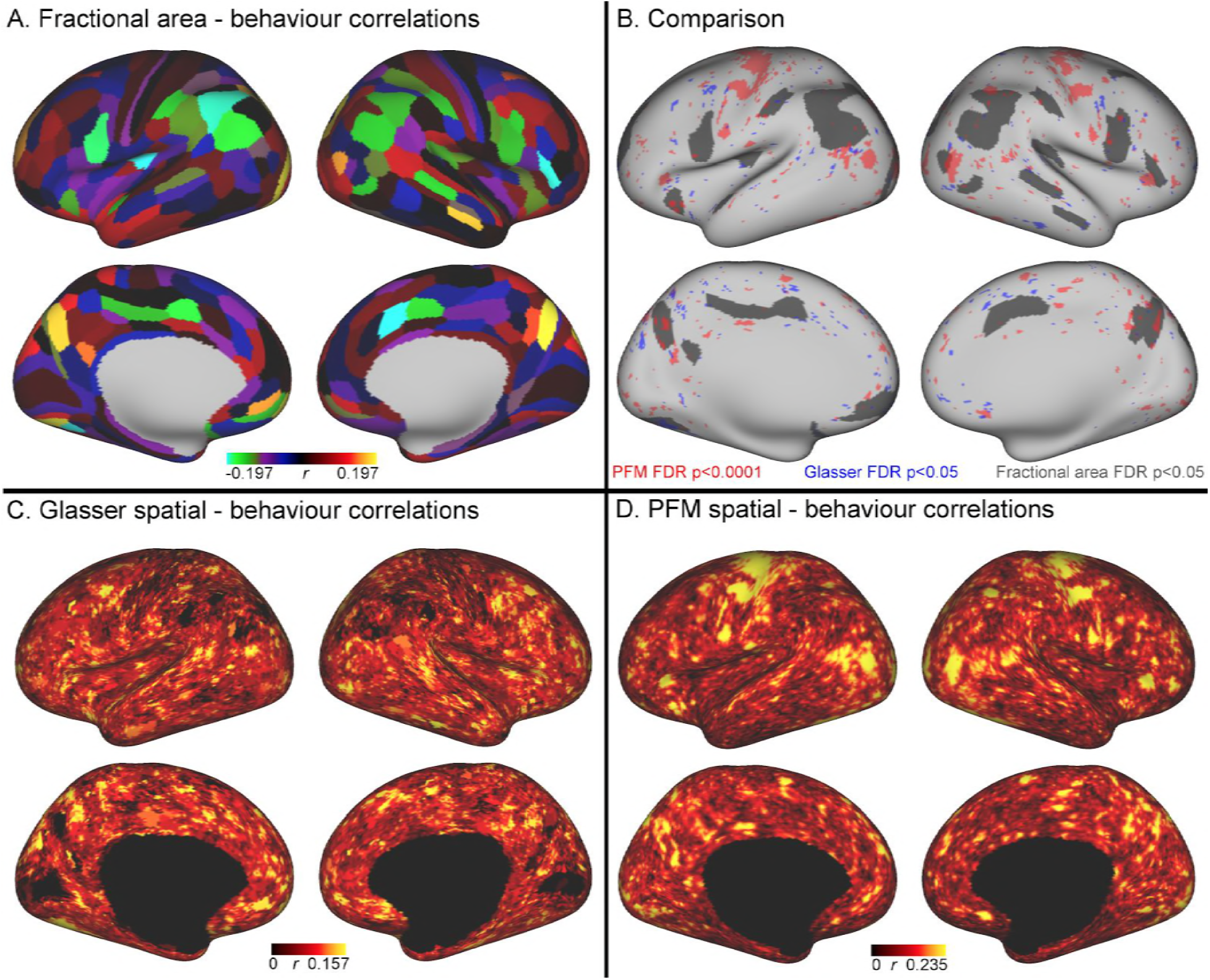
Comparison of the cortical representation of associations with behaviour across fractional area, HCP_MMP1.0 individual subject parcellation and PFM spatial maps. A: Correlations between fractional surface area and behaviour are bilaterally symmetric and strongest in cognitive and association cortices. B: Direct comparison of strongest results qualitatively suggests that PFM and HCP_MMP1.0 effects may be partly linked to areal size, and highlights lack of precise co-location of effects (for visual comparison the PFM correlation maps are shown using a higher threshold p_FDR_<0.0001, |r|>0.218, and HCP_MMP1.0 correlation maps are correlated at p_FDR_<0.05; |r|>0.159). C: Un-thresholded HCP_MMP1.0 correlations with CCA subject weights; these are the maximum absolute r across all parcels, and therefore do not contain the parcel structure itself. D: Un-thresholded PFM correlations with CCA subject weights (maximum absolute r across all PFMs).

### Spatiotemporal simulations demonstrating potential sources of variability in edges

The ICA 200 full network matrix results shown in the main manuscript Table 1 reveal that highly similar network matrices can be obtained from simulated data for which the only source of cross-subject variability is differences in spatial maps. We considered whether these results might be specific to the timeseries extraction approach used to create network matrices (for Table 1 this was dual regression against ICA 200 dimensional group maps). To address this, we repeated the same analysis (using the identical approach to generate simulated datasets), performing dual regression against a lower dimensional set of group maps (d=25), and also after masking against two different types of parcellations (HCP_MMP1.0 & Yeo). For the HCP_MMP1.0 parcellation, parcel definitions are available both at the group and subject level, and both were tested here. The results (Table S3) for a wide range of different parcellations show comparable trends (i.e., a large proportion of cross-subject variability is captured purely by spatial maps, as indicated by the highlighted rows).

**Table S3:** Results from simulated datasets in which one or more of the network matrices, amplitudes and spatial maps are fixed to the group average to remove any subject variability associated with it. Results in each row were driven by variables in which subject variability was present, as indicated with ✓ (variables with - were fixed to the group average). Results are shown for within-subject correlations between simulated and original z-transformed network matrices (Z_network matrix_), across-subject correlations between simulated and original subject correlation matrices (R_correlation_), and for results obtained from the CCA against behaviour. Note that comparable CCA results from the original data can be found in Table S1.This Table presents results from full correlation network matrices.

| Full network matrices | Results driven by subject variability in: | Network matrix | Amplitudes | Spatial maps | Z network matrix | R correlation | CCA $r_{U-V}$ | CCA $P_{U-V}$ | CCA $r_{U-Uica}$ |
| --- | --- | --- | --- | --- | --- | --- | --- | --- | --- |
| ICA<br>D = 200<br>N=819 | Nothing | - | - | - | -0.0003 | 0.03 | 0.65 | 0.32017 | 0.11 |
|  | Amps & spatial | - | ✓ | ✓ | 1.14 | 0.60 | 0.71 | 0.00001 | 0.52 |
|  | Connectivity only | ✓ | - | - | 0.47 | 0.65 | 0.69 | 0.00028 | 0.40 |
|  | Amplitudes only | - | ✓ | - | 0.22 | 0.15 | 0.69 | 0.00052 | 0.45 |
|  | Spatial maps only | - | - | ✓ | 0.78 | 0.54 | 0.72 | 0.00001 | 0.62 |
| ICA<br>D = 25<br>N=819 | Nothing | - | - | - | -0.0004 | -0.003 | 0.65 | 0.26790 | 0.12 |
|  | Amps & spatial | - | ✓ | ✓ | 1.19 | 0.52 | 0.71 | 0.00001 | 0.47 |
|  | Connectivity only | ✓ | - | - | 0.88 | 0.75 | 0.69 | 0.00005 | 0.44 |
|  | Amplitudes only | - | ✓ | - | 0.26 | 0.08 | 0.67 | 0.03876 | 0.47 |
|  | Spatial maps only | - | - | ✓ | 0.78 | 0.61 | 0.73 | 0.00001 | 0.47 |
| Yeo | Nothing | - | - | - | -0.002 | -0.01 | 0.65 | 0.14899 | 0.14 |
| parcellation<br>D = 109<br>N=819 | Amps & spatial | - | ✓ | ✓ | 1.09 | 0.40 | 0.72 | 0.00001 | 0.59 |
|  | Connectivity only | ✓ | - | - | 0.50 | 0.55 | 0.69 | 0.00003 | 0.47 |
|  | Amplitudes only | - | ✓ | - | 0.25 | 0.10 | 0.68 | 0.00258 | 0.27 |
|  | Spatial maps only | - | - | ✓ | 0.69 | 0.40 | 0.74 | 0.00001 | 0.61 |
| HCP_MMP<br>1.0 subject<br>parcellation<br>D = 360<br>N=441 | Nothing | - | - | - | 0.19 | 0.19 | 0.85 | 0.02565 | 0.34 |
|  | Amps & spatial | - | ✓ | ✓ | 1.04 | 0.38 | 0.86 | 0.00055 | 0.51 |
|  | Connectivity only | ✓ | - | - | 0.46 | 0.62 | 0.85 | 0.01399 | 0.27 |
|  | Amplitudes only | - | ✓ | - | 0.30 | 0.14 | 0.84 | 0.17321 | 0.34 |
|  | Spatial maps only | - | - | ✓ | 0.65 | 0.45 | 0.85 | 0.01161 | 0.70 |
| HCP_MMP<br>1.0 group<br>parcellation<br>D = 360<br>N = 441 | Nothing | - | - | - | -0.0005 | -0.01 | 0.85 | 0.01726 | 0.18 |
|  | Amps & spatial | - | ✓ | ✓ | 1.03 | 0.41 | 0.86 | 0.00153 | 0.48 |
|  | Connectivity only | ✓ | - | - | 0.43 | 0.62 | 0.86 | 0.00034 | 0.37 |
|  | Amplitudes only | - | ✓ | - | 0.21 | 0.11 | 0.84 | 0.18253 | 0.29 |
|  | Spatial maps only | - | - | ✓ | 0.65 | 0.43 | 0.86 | 0.00850 | 0.61 |

Partial network matrices may be better suited to control for misalignment than full network matrices. Therefore, we repeated to full set of analyses presented in Figure 1 and Table S3 after estimating partial network matrices from the timeseries extracted from simulated data. The results (Table S4) show that the main result is also found when using partial network matrices (e.g., for ICA 200, 0.51^2^=26% variance explained in partial network matrices was captured by spatial information, and 0.54^2^=29% variance explained in full network matrices was captured by spatial information). Therefore, partial network matrices are also strongly influenced by cross-subject spatial variability.

Note that the results driven by subject network matrices for partial correlations of ICA 200 dimensionality, and the Yeo and HCP_MMP1.0 parcellation in Table S4 are considerably lower compared with the full correlation findings for the same simulation (Table S3). The reason for this is that the PFM 50-dimensional subject network matrices were added into the data (to keep the simulation pipeline identical). This approximated 50-dimensional network matrix is too low rank to allow accurate estimation of partial connectivity across a much larger number of nodes. The full correlation results in Table S3 are estimable, and support the 25-dimensional ICA results.

**Table S4:** Results from simulated datasets in which one or more of the network matrices, amplitudes and spatial maps are fixed to the group average to remove any subject variability associated with it. Results in each row were driven by variables in which subject variability was present, as indicated with ✓ (variables with - were fixed to the group average). Results are shown for within-subject correlations between simulated and original z-transformed network matrices (Z_network matrix_), across-subject correlations between simulated and original subject correlation matrices (R_correlation_), and for results obtained from the CCA against behaviour. This Table presents results from partial correlation network matrices. Note that the results flagged with * are poorly estimated as a result of the low rank of the PFM subject network matrices (containing 50 PFM modes) used to drive these simulations.

| Partial network matrices | Results driven by subject variability in: | Network matrix | Amplitudes | Spatial maps | Z network matrix | R correlation | CC A r U-V | CCA P U-V | CCA r U-Uica |
| --- | --- | --- | --- | --- | --- | --- | --- | --- | --- |
| ICA<br>D = 200<br>N=819 | Nothing | - | - | - | -0.0003 | 0.02 | 0.65 | 0.18364 | 0.15 |
|  | Amps & spatial | - | ✓ | ✓ | 0.61 | 0.58 | 0.77 | 0.00001 | 0.78 |
|  | Connectivity only | ✓ | - | - | 0.02* | 0.06* | 0.66* | 0.0417* | 0.14* |
|  | Amplitudes only | - | ✓ | - | 0.08 | 0.12 | 0.68 | 0.00109 | 0.42 |
|  | Spatial maps only | - | - | ✓ | 0.53 | 0.51 | 0.76 | 0.00001 | 0.77 |
| ICA<br>D = 25<br>N=819 | Nothing | - | - | - | -0.001 | -0.02 | 0.64 | 0.48333 | 0.11 |
|  | Amps & spatial | - | ✓ | ✓ | 0.96 | 0.50 | 0.71 | 0.00001 | 0.54 |
|  | Connectivity only | ✓ | - | - | 0.56 | 0.38 | 0.69 | 0.00007 | 0.24 |
|  | Amplitudes only | - | ✓ | - | 0.15 | 0.11 | 0.67 | 0.03009 | 0.39 |
|  | Spatial maps only | - | - | ✓ | 0.73 | 0.65 | 0.70 | 0.00001 | 0.50 |
| Yeo parcellation<br>D = 109<br>N=819 | Nothing | - | - | - | 0.0007 | 0.02 | 0.64 | 0.57151 | 0.15 |
|  | Amps & spatial | - | ✓ | ✓ | 0.91 | 0.29 | 0.73 | 0.00001 | 0.53 |
|  | Connectivity only | ✓ | - | - | 0.09* | 0.29* | 0.67* | 0.0192* | 0.35* |
|  | Amplitudes only | - | ✓ | - | 0.11 | 0.001 | 0.68 | 0.00312 | 0.39 |
|  | Spatial maps only | - | - | ✓ | 0.75 | 0.72 | 0.76 | 0.00001 | 0.63 |
| HCP_MMP 1.0 subject parcellation<br>D = 360<br>N=441 | Nothing | - | - | - | 0.27 | 0.40 | 0.85 | 0.04779 | 0.36 |
|  | Amps & spatial | - | ✓ | ✓ | 0.81 | 0.36 | 0.87 | 0.00038 | 0.58 |
|  | Connectivity only | ✓ | - | - | 0.30* | 0.44* | 0.86* | 0.0064* | 0.37* |
|  | Amplitudes only | - | ✓ | - | 0.25 | 0.03 | 0.85 | 0.06235 | 0.46 |
|  | Spatial maps only | - | - | ✓ | 0.73 | 0.79 | 0.88 | 0.00001 | 0.55 |
| HCP_MMP 1.0 group parcellation<br>D = 360<br>N = 441 | Nothing | - | - | - | -0.0003 | -0.05 | 0.84 | 0.11041 | 0.17 |
|  | Amps & spatial | - | ✓ | ✓ | 0.76 | 0.33 | 0.87 | 0.00018 | 0.60 |
|  | Connectivity only | ✓ | - | - | 0.07* | 0.36* | 0.85* | 0.0491* | 0.20* |
|  | Amplitudes only | - | ✓ | - | 0.05 | -0.02 | 0.84 | 0.22268 | 0.26 |
|  | Spatial maps only | - | - | ✓ | 0.68 | 0.74 | 0.87 | 0.00005 | 0.59 |

### Role of spatially varying amplitudes

The subject-specific PFM maps obtained from PROFUMO are relatively complex and can represent a range of different types of information, including the shape and size of the modes, but also the relative amplitudes of separate network nodes included in the PFM spatial modes. The latter might be classed as information that represents (within-mode) functional connectivity, rather than being a purely spatial feature of the mode. We performed a further set of simulations in which we aimed to separate the role of subject-specific spatial features (such as shape and size) from the role of these complex amplitudes included in the spatial map. In order to do this, we thresholded and binarized the PFM spatial maps prior to generating the simulated datasets, in order to simplify the maps and remove the amplitude-information contained in them. This binarization procedure was performed using either a fixed threshold across subjects (which may allow differences in the size of resulting maps across subjects), or using a percentile threshold (calculated within subject and mode) to fix the size of the binarized maps across subjects and only allow shape information to drive the simulation. Therefore, the cross-subject variability present in these simulated datasets was purely driven by the spatial shape of functional networks (amplitude and network matrices were both fixed to the group average). The results of network matrices extracted from these binarized-map-based simulations using either dual regression against ICA-200 maps or masking against Yeo parcellations are presented in Table S5 and show that the role of the spatial maps in driving the resulting functional connectivity estimates remains comparably strong (for example, all CCA results in table S5 are highly significant; p=0.00001), even when highly simplified maps that only contain a binarized representation of shape are used.

**Table S5:** Modulating the subject spatial maps by thresholding and binarizing retains the shape and size aspects, but removes any relative amplitude information from the spatial maps. Binarized % results are binarized after applying a percentile threshold, and therefore only retain shape aspects (while fixing the size). The results reveal that even after thresholding and binarizing the spatial maps, remaining spatial variability strongly drives the cross-subject information present in the resulting network matrices. See earlier Tables for a description of the measures.

|  | **Spatial maps** | **Z<sub>network matrix</sub>** | **R<sub>correlation</sub>** | **CCA r<sub>U-V</sub>** | **CCA P<sub>U-V</sub>** | **CCA r<sub>U-Uica</sub>** |
| --- | --- | --- | --- | --- | --- | --- |
| ICA 200 Full | Thresholded | 0.43 | 0.32 | 0.74 | 0.00001 | 0.67 |
|  | Binarized | 0.45 | 0.43 | 0.74 | 0.00001 | 0.63 |
|  | Binarized % | 0.44 | 0.44 | 0.70 | 0.00001 | 0.57 |
| ICA 200 Partial | Thresholded | 0.29 | 0.42 | 0.74 | 0.00001 | 0.73 |
|  | Binarized | 0.28 | 0.54 | 0.76 | 0.00001 | 0.77 |
|  | Binarized % | 0.41 | 0.55 | 0.77 | 0.00001 | 0.76 |
| Yeo 109 Full | Thresholded | 0.57 | 0.37 | 0.71 | 0.00001 | 0.56 |
|  | Binarized | 0.56 | 0.37 | 0.71 | 0.00001 | 0.54 |
|  | Binarized % | 0.44 | 0.37 | 0.72 | 0.00001 | 0.56 |
| Yeo 109 Partial | Thresholded | 0.48 | 0.68 | 0.75 | 0.00001 | 0.63 |
|  | Binarized | 0.46 | 0.62 | 0.75 | 0.00001 | 0.67 |
|  | Binarized % | 0.54 | 0.62 | 0.75 | 0.00001 | 0.52 |

### Comparing cross-subject similarities between different types of imaging measures

The simulation results and CCA results presented above suggest that the same cross-subject variance structure is present in a range of different rfMRI-derived measures. Given that cross-subject variability is typically of key interest in neuroimaging studies, the next analysis aimed to directly compare the subject variations present in the network matrices, spatial maps and amplitudes obtained from the original data. This is achieved by calculating subject-by-subject correlation matrices from different measures and comparing these matrices between the imaging measures.

The results show that the cross-subject variability that is present in estimated network matrices is shared with ICA spatial maps, PFM spatial maps and PFM amplitudes (Figure S2). Of these, subject variations in estimated network matrices are most similar to PFM spatial maps, and these similarities remain relatively strong when looking at the partial correlations (i.e., after regressing out any variance that can be explained by the other factors). These findings are consistent with the simulation results above, showing that estimated network matrices and spatial topography to a large extent overlap in terms of the interesting cross-subject variability they represent. Additionally, the results show that while dual regression ICA spatial maps are able to capture some of the subject spatial variability, subject maps estimated by PROFUMO capture considerably more spatial variability over and above the dual regression maps.

**Figure S2:**
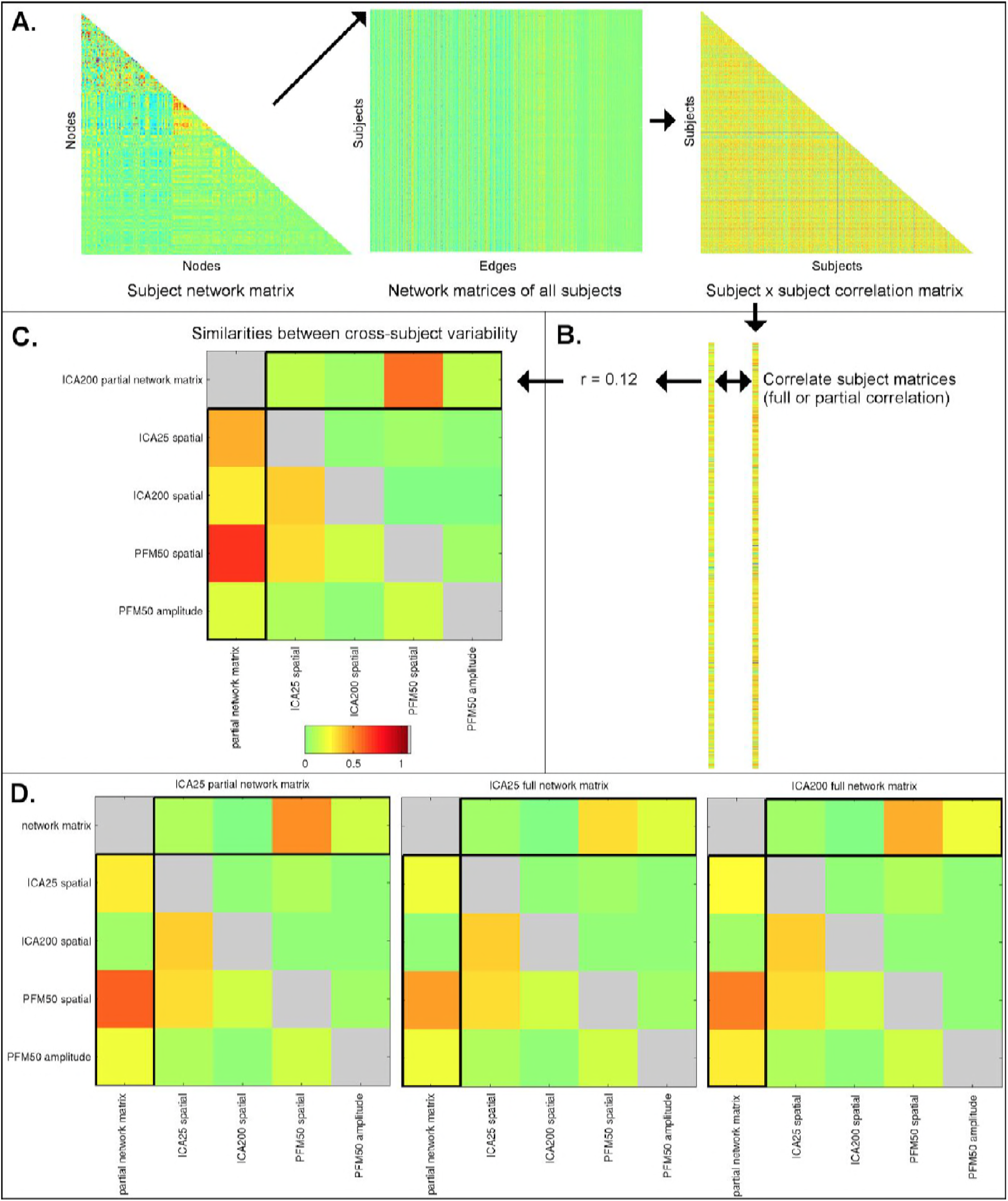
Similarities between cross-subject variations estimated from different rfMRI measures. Subject-by-subject correlation matrices are estimated (A), and vectorised (B; one subject correlation matrix being estimated for each measure type). The first column of the similarities (C; highlighted) shows the relationship (full correlation) between the ICA network matrix and various other measures, such as PFM spatial maps and amplitudes, and ICA spatial maps. These results show that the ICA network matrix is closely related to PFM spatial maps. The first row of the similarities (C; highlighted) shows the same relationship after taking into account all the other elements (i.e., the partial correlation between different measures). This reveals that PFM spatial maps are strongly linked to the ICA network matrix, even after accounting for any variance that can be explained by ICA spatial maps and PFM amplitudes. Similar results are obtained for ICA 200 and 25 dimensionality and for partial and full network matrices (D).

